# In quiescent G0 phase, *Schizosaccharomyces pombe* Mis4 ensures full nuclear separation during the subsequent M phase

**DOI:** 10.1101/2024.03.29.587322

**Authors:** Michiko Suma, Orie Arakawa, Yuria Tahara, Kenichi Sajiki, Shigeaki Saitoh, Mitsuhiro Yanagida

**Author notes:** **Corresponding authors:** S. Saitoh, M. Yanagida.

## Abstract

Evolutionarily conserved Mis4 establishes cohesion between replicated sister chromatids in vegetatively proliferating cells. In the fission yeast, *Schizosaccharomyces pombe*, defects in Mis4 lead to premature separation of sister chromatids, resulting in fatal chromosome mis-segregation during mitosis. In humans, NIPBL, an ortholog of Mis4, is responsible for a severe multisystem disorder called Cornelia de Lange syndrome. We previously reported that Mis4 is also essential in non-proliferating quiescent cells. Whereas wild-type fission yeast cells can maintain high viability for long periods without cell division in the quiescent G0 phase, *mis4-450* mutant cells cannot. In this report, we show that Mis4 is not required for cells to enter G0 phase, but is essential to exit from it. When resuming mitosis after passage of G0, *mis4* mutant cells segregated sister chromatid successfully, but insufficiently separated daughter nuclei consequently formed dikaryon-like cells. These findings suggest a novel role of Mis4/NIPBL in non-dividing quiescent cells, which is prerequisite for full nuclear separation upon resumed mitosis. As most human cells are in the quiescent state, this study may facilitate development of novel therapies for human diseases caused by Mis4/NIPBL deficiency.

## Introduction

In eukaryotes, precise segregation of sister chromatids to daughter cells is essential to maintain genome integrity and cell viability. Segregation failure results in cell death and disease in humans. Cohesin, a ring-like protein complex, holds sister chromatids together until the onset of mitotic anaphase (Biggins and Murray,1998; Gruber and Haering, 2003). Fission yeast Mis4 reportedly functions as a cohesin loader, mounting the complex on chromatin (Murayama and Uhlmann, 2014). Mis4 is thought to function specifically during S phase, as its temperature-sensitive (*ts*) mutation causes lethal mis-segregation of sister chromatids during mitosis after passing S phase at restrictive temperatures (Furuya et al., 1998). Mis4 is conserved evolutionarily from yeasts to humans and its orthologues are designated as Nipped-B and Scc2 in *Drosophila melanogaster* and budding yeast (*Sacchromyces cerevisiae*), respectively (Moore et al., 1998; Rollins et al., 1999; Tó et al., 1999). These orthologues also function in gene expression, gene silencing, and DNA damage repair (Donze et al., 1999; Zakari et al., 2015). In budding yeast, mutant cells defective in Scc2 shows ectopic expansion of silenced chromatin, which is often accompanied by frameshift mutations. In humans, NIPBL, the Mis4 orthologue, is related to severe diseases, such as Cornelia de Lange syndrome (CdLS) and breast cancer (Tonkin et al., 2004; Zhou et al., 2017). CdLS, the prevalence of which is estimated to be 1 in 10,000, is a multisystem disorder accompanied by growth and cognitive retardation. In human cell lines derived from breast cancer, MFC7 and Bcap37, the NIPBL gene and protein expression are upregulated, and downregulation of NIPBL expression results in cell cycle arrest in the G1/G0 phase and induced apoptosis (Zhou et al., 2017).

In multicellular organisms, most differentiated cells are in the quiescent G0 phase, during which they are metabolically active, but stop dividing (Mochida and Yanagida, 2006). Failure to enter quiescence is thought to cause excessive cell proliferation and consequent tumorigenesis. Switching between vegetative proliferation and quiescence is regulated by various stimuli, such as depletion/repletion of nutrients and growth factors (Pardee, 1974). In fission yeast, depletion of nitrogen sources, such as ammonium chloride, from the medium prompts cells to enter the G0 phase, from which they recover to the vegetatively proliferating state upon replenishment of nitrogen (Su et al., 1996). Previously, we have screened libraries of fission yeast *ts* mutant strains and gene deletion strains for those that lose cell viability under nitrogen-depletion (Sajiki et al., 2009; Sajiki et al., 2018). Responsible genes were identified and collectively designated as *shk* (*s*uper *h*ouse *k*eeping) and *gze* (*g*-*z*ero *e*ssential) genes, respectively. *shk* genes are apparently required for maintenance of cell viability in both proliferating and quiescent cells, whereas *gze* genes are essential only in quiescent cells. 40 *shk* genes and 85 *gze* genes have been so far identified (Hayashi et al., 2018; Nakazawa et al., 2018; Sajiki et al., 2009; Sajiki et al., 2018; Uehara et al., 2021). As diverse molecular roles of these gene products are predicted from annotations in POMBASE (https://www.pombase.org), a wide variety of molecules function together to enable cells to enter or exit G0 phase. Notably, *mis4* has been identified as a *shk* gene (Sajiki et al., 2009), implying that Mis4 is essential in quiescent cells; however, its exact molecular function remains unknown, as do its activities in sister chromatid cohesion in vegetatively proliferating cells.

In this study, to determine the physiological roles of Mis4 in non-dividing, quiescent cells, cell cycle progression and mitotic nuclear division were examined in *mis4* mutant cells upon recovery from G0 quiescence. Whereas chromosome segregation was apparently normal in these mutant cells, subsequent nuclear separation proved defective, resulting in accumulation of abnormal dikaryon-like cells that contained two adjacent, interphase nuclei in single cell bodies. These findings suggest a novel role of evolutionarily conserved Mis4 in non-dividing, quiescent cells that ensures proper spindle elongation and/or nuclear separation upon resuming vegetative cell proliferation. This study may help to clarify molecular mechanisms by which NIPBL/Mis4 deficiency causes various symptoms of CdLS and breast cancer.

## Results

### The fission yeast *mis4-450* mutant loses viability under nitrogen starvation

*Schizosaccharomyces pombe* cells harboring the *mis4-450* mutation show the *shk* phenotype, losing viability under both vegetative and quiescent conditions at restrictive and/or semi-permissive temperatures (Sajiki et al., 2009). To identify mutation sites in the *mis4-450* mutant allele, a DNA fragment containing the *mis4* gene was amplified from the mutant genome with high-fidelity PCR, and its sequence was determined. It contains two missense mutations causing amino acid substitutions, L296F and P368L (**Fig. 1A**). To confirm that these mutations are responsible for the temperature sensitivity of the *mis4-450* mutant, strains harboring one or both were generated by DNA recombination and their colony formation abilities were examined on nutrient-rich YPD medium at various temperatures. *mis4* DNA fragments containing L296F, P368L, or both were introduced to wild-type cells (strain: 972 *h*^-^), and cells in which the authentic *mis4*^+^ gene was replaced with the introduced mutant *mis4* gene were selected (**Fig. 1B**). Resulting strains, which were designated M162, M165, and M166, were used in further experiments in this study. For a wild-type control (WT), a DNA fragment containing the wild-type allele of *mis4* was introduced into wild-type cells, and the resulting strain was designated M161. Strains with either L296F or P368L alone (M165 and M166) grew as well as the WT (M161) at all temperatures examined, whereas the strain harboring both mutations together (M162) failed to form colonies at 36 °C (**Fig. 1C**), indicating that the combination of these two missense mutations in the *mis4* gene causes the *ts* phenotype.

**Figure 1.**
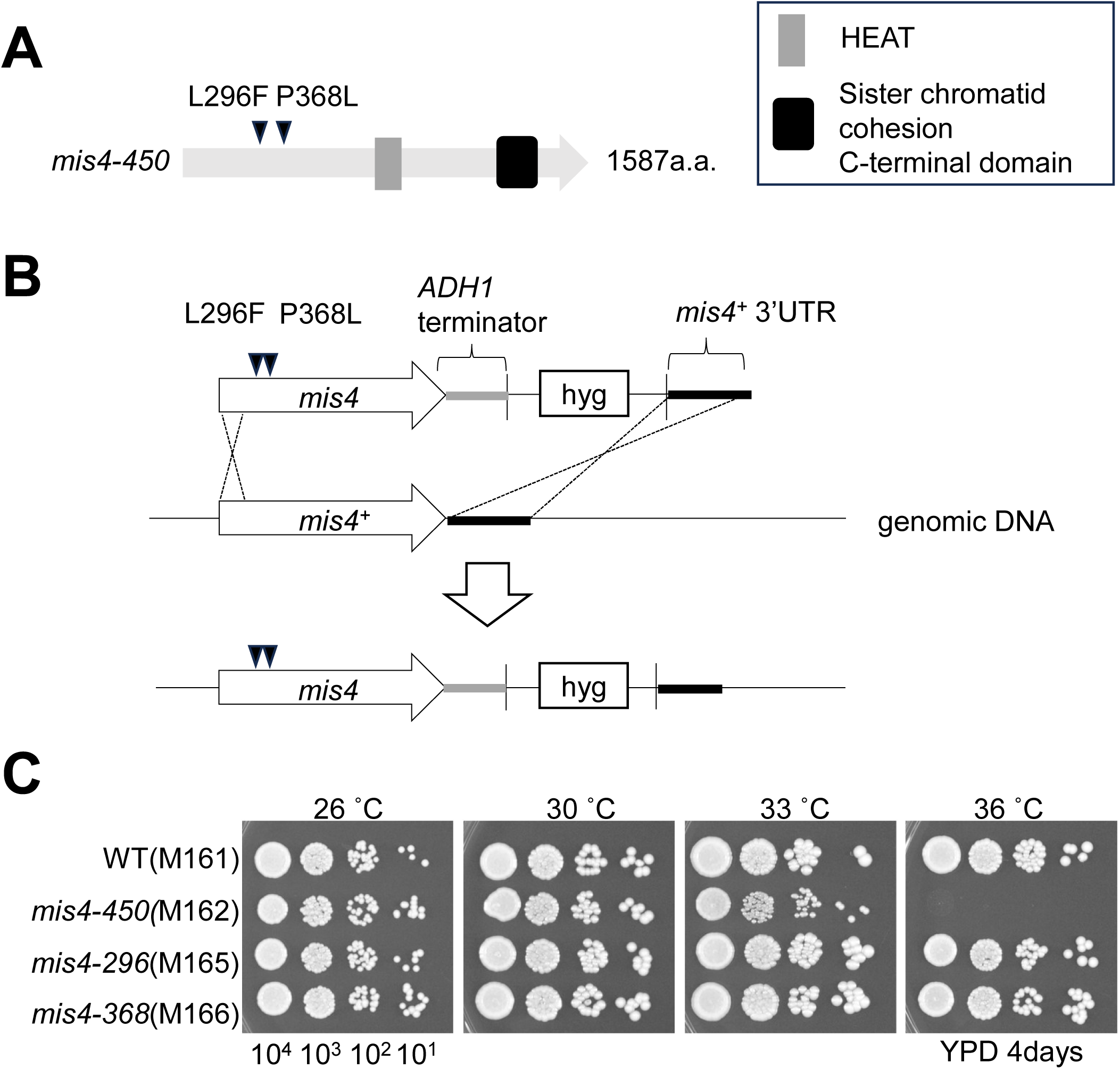
Two amino acids, L296 and P368, are mutated in the *mis4-450 ts* mutant allele. **(A)** Positions of missense mutations in *mis4-450* were determined by nucleotide sequencing, and schematically shown with characteristic domains of Mis4/Scc2/Nipped-B family proteins. *S. pombe* Mis4 contains a HEAT repeat (801 – 842 aa, gray box) and sister chromatid cohesin C-terminal domain (1301 – 1482 aa, black box). Mutation sites of the *mis4-450* allele are not located in these domains. **(B)** To generate strains M162, M165, and M166, the *mis4* gene in *S. pombe* wild-type cells (strain: 972 *h*^-^) was replaced with *mis4* DNA fragments containing L296F, P368L or both mutations, by homologous recombination. See *Materials and Methods* for details. **(C)** Aliquots of 10^4^ cells of WT control (M161), M162, M165, and M166 strains were serially diluted 10-fold, spotted onto YPD solid medium and incubated at 26 ℃, 30 ℃, 33 ℃ and 36 ℃ for 4 days. Only M162 cells, which harbor both L296F and P368L mutations in *mis4*, failed to form colonies at 36 ℃, indicating that the combination of these mutations confers temperature sensitivity.

We then examined whether the L296F and P368L mutations in the *mis4* gene cause the *shk* phenotype, cells that lose cell viability after cultivation in nitrogen-depleted medium prompting them to enter the G0 phase. Cells with the L296F and P368L mutations in *mis4* (M162) and WT cells (M161) were first cultivated in liquid EMM2 medium containing 93.5 mM ammonium chloride, and were then transferred to EMM2 lacking ammonium chloride (EMM2-N). Aliquots of the cell culture were spread on solid YPD medium to measure cell viability as determined from the proportion of cells forming colonies under permissive conditions (**Fig. 2A**). Consistent with a previous report (Sajiki et al., 2009), the viability of M162 cells decreased to 22% after 22 day-incubation in EMM2-N medium at 26 °C, while that of WT cells remained as high as ∼70%, indicating that these mutations in the *mis4* gene are responsible for the *shk* phenotype. Importantly, viability of the mutant greatly decreased even though cells were cultivated at 26 °C, a permissive temperature for *mis4* mutant cells when incubated on nutrient-rich medium (**Fig. 1C**), suggesting that even a slight malfunction of Mis4, which may occur at 26 °C and which can be tolerated for mitotic cell division, leads to significant reduction of cell viability in non-dividing quiescent cells.

**Figure 2.**
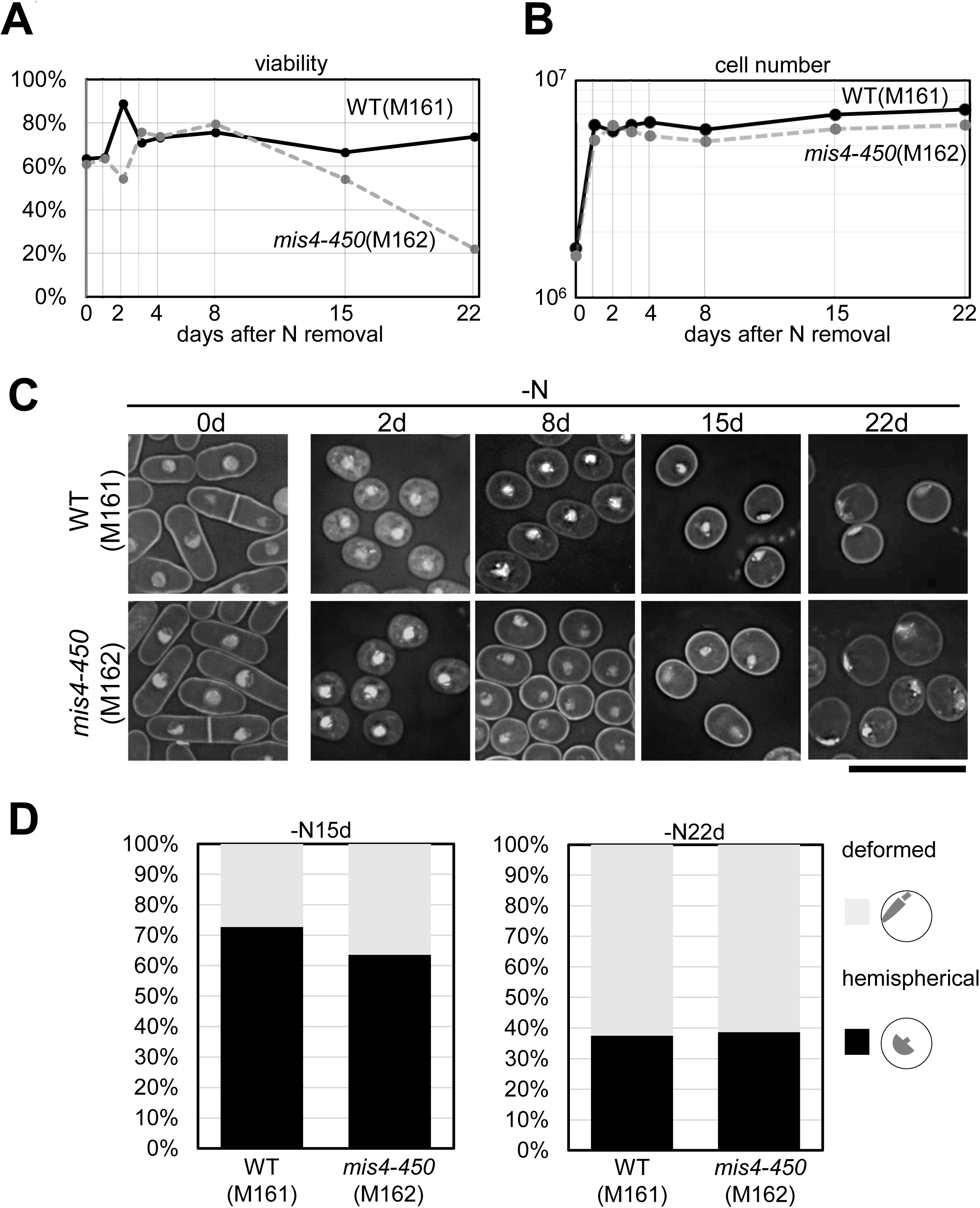
The *mis4-450* mutation reduces cell viability after 22-day nitrogen starvation. **(A)** Cell viability under nitrogen-depleted conditions. WT control (M161 strain, solid line) and the *mis4-450* mutant (M162 strain, dotted line) cells were transferred from nitrogen-rich EMM2 (93.4mM NH_4_Cl) to nitrogen-depleted EMM2 (EMM2-N) medium and cultivated for 22 days. Small aliquots of cell culture were taken to measure cell viability at the indicated time points after the transfer to nitrogen-depleted medium. **(B)** Concentrations of WT control (M161, solid line) and *mis4-450* mutant (M162, dotted line) cells transferred to nitrogen-depleted medium were measured at the indicated time points. Both WT (M161) and *mis4* mutant (M162) cells divided twice to increase their concentrations by ∼4 times within 1 day after the transfer. WT (M161) and the mutant (M162) cells stopped increasing their numbers thereafter. **(C)** Morphologies of cells and nuclei were examined in WT and *mis4* mutant cells grown in nitrogen-depleted medium. WT (M161) and *mis4-450* mutant (M162) cells transferred from nitrogen-rich EMM2 medium to nitrogen-depleted medium (-N) were fixed with 10% glutaraldehyde before the transfer (0d) and at 2 days (2d), 8 days (8d), 15 days (15d) and 22 days (22d) after the transfer. Then they were stained with DAPI. Three-dimensional Z-serial fluorescent images, which were taken using a Delta Vision microscope, were deconvoluted and flattened using SoftWoRx imaging workstation. Bar, 10μm. **(D)** Frequencies of cells with hemispherical nuclei (black) and those with deformed “comet-shaped” (Su et al., 1996) nucleus (gray) are shown. 200 cells after 15-day (left panel) and 22-day (right panel) nitrogen starvation, were examined to determine frequencies. In both WT (M161) and *mis4* mutant (M162) strains, the frequency of cells with hemispherical nuclei decreased to ∼40% and those with comet-shaped nucleus increased to ∼60% 22 days after the transfer to EMM2-N medium.

Changes in cell number and morphology after the shift to EMM2-N medium are shown in **Figs. 2B** and **2C**. As with WT control cells, *mis4* mutant cells divided twice and their number increased by ∼4 times before ceasing cell proliferation in EMM2-N medium (**Fig. 2B**). Changes in shapes of cells and nuclei were identical in WT controls and *mis4* mutants. Cells became small and round 1 day after the medium shift, and nuclei were deformed near the cell surface after incubation in -N medium for > 15 days (**Fig. 2C and D**). These changes are characteristic of G0 quiescent cells (Su et al, 1996); thus, it is apparent that the L296F and P368L mutations in the *mis4* gene do not prevent cells from entering G0 phase.

Notably, while four *ts* mutant alleles of the *mis4* gene (*mis4-242, -236*, *-381* and *-450*) were found in our *ts* mutant library and all of them cause abnormal chromosome segregation due to defective chromatid cohesion during mitosis, only the *mis4-450* mutant allele caused the *shk* phenotype (Sajiki et al., 2009) (**Supplemental Fig. S1A and B)**. The *mis4-242*, *mis4-236*, and *mis4-381* alleles were found to harbor single, missense mutations: G1326E, F508S and S1314P, respectively (**Supplemental Fig. S1C**). Given that the mutation sites of the *mis4-450* allele are distant from those of the other alleles, distinct domains of Mis4 protein may function separately in chromatid cohesion and cellular quiescence.

### *mis4-450* mutation causes insufficient nuclear separation when cells leave G0 quiescence to resume mitotic cell cycle progression

As shown above, M162 cells, which harbor L296F and P368L mutations in the *mis4* gene, lose cell viability during G0 quiescence, although their entry into G0 appears normal. These results raise the possibility that the *mis4-450* mutant cannot resume mitotic cell cycle properly after being transferred from G0-inducing, nitrogen-depleted medium to that with sufficient nitrogen. To test this possibility, mitotic cell cycle progression was examined in *mis4* mutant (M162) and WT control cells, which were incubated in EMM2-N medium for 22 days at 26 °C and then transferred to nutrient-rich YPD complete medium (**Fig. 3A**). Fluorescence-activated cell sorting analysis (FACScan) of the DNA content revealed that completion of DNA synthesis was delayed slightly in *mis4* mutant cells (**Fig. 3B)**. Most WT cells contained replicated 2C DNA 12 h after the shift to YPD, whereas a significant portion of mutant cells contained unreplicated 1C DNA at the same time point. Intriguingly, nuclear separation following mitotic chromosome segregation appeared insufficient in *mis4* mutant cells. Cells containing two interphase nuclei, located close to each other near the cell center, were frequently observed in M162 cells after the shift to the YPD medium (**Fig. 3C-3D**), while such cells were rarely seen in WT controls. This phenotype resembles a dikaryon phenotype previously reported (Grallert et al., 1998; Okazaki and Niwa, 2008). The frequency of such cells started to increase 18 h after the shift to YPD medium in *mis4* mutants (M162), and reached ∼30% of binucleated cells at 20 h. Importantly, the insufficient nuclear separation phenotype described above arose only after passage of the G0 quiescence phase. *mis4* mutant cells that were cultivated continuously in YPD medium at 26 °C did not exhibit such a phenotype, and appeared identical to WT control cells (**Fig.4A and B**).

**Figure 3.**
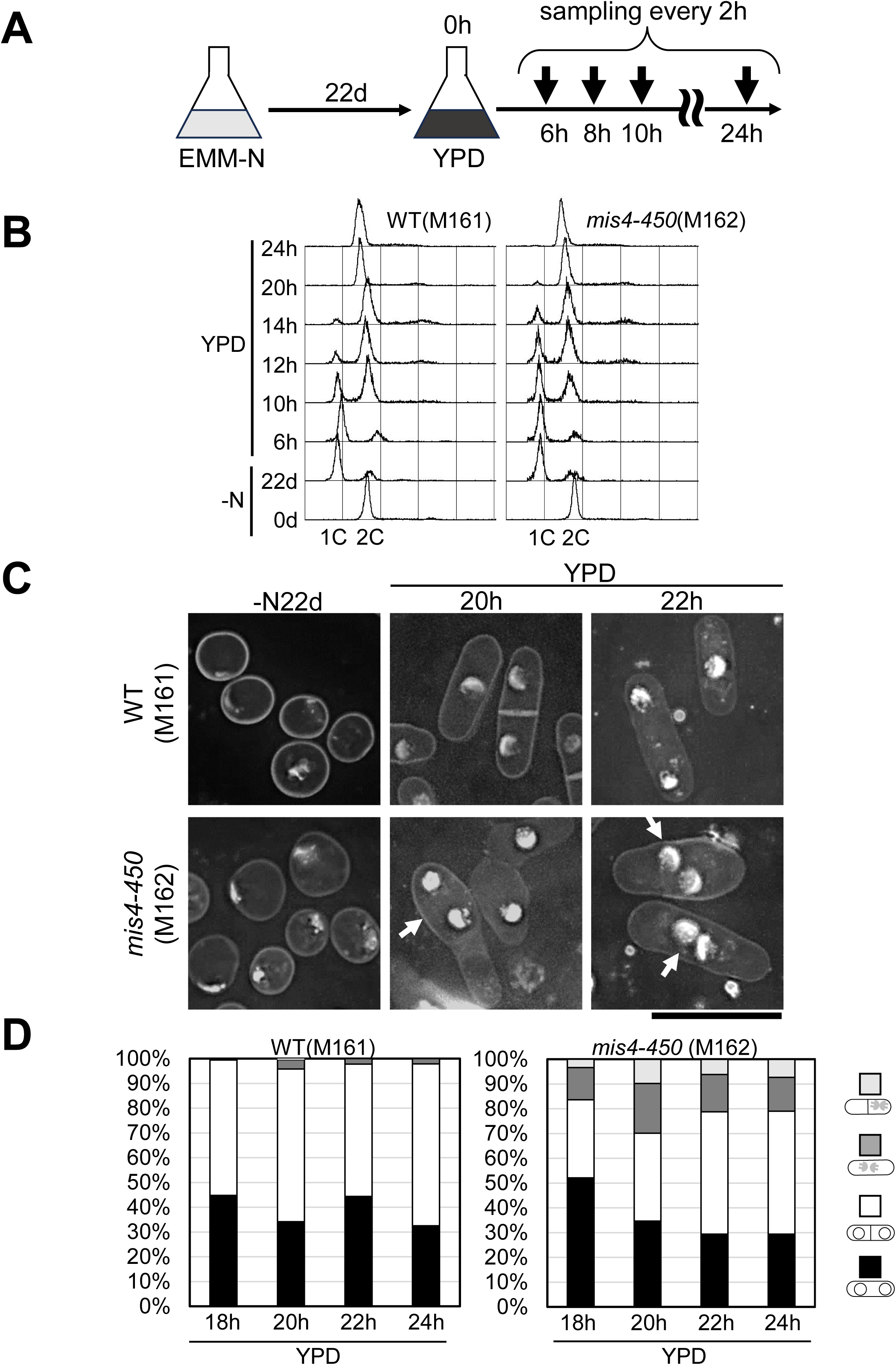
*mis4-450* mutant cells exhibit a dikaryon-like phenotype during resumption of mitotic cell division after 22-day nitrogen starvation. **(A)** The experimental procedure for recovery from G0 quiescence is shown. Cells proliferating vegetatively in nitrogen-rich YPD medium were transferred to EMM2-N medium for entry into G0. After 22-day cultivation in EMM2-N medium, during which cells remained in G0 quiescence, cells were transferred to YPD medium for re-entry into the mitotic cell cycle. Aliquots of the cell culture were taken at 2-h intervals for analyses of DNA content (B) and analyses of cell morphology and nuclei (C and D). **(B)** DNA contents were measured by flow cytometry in WT (M161) and *mis4-450* (M162) mutant cells after the transfer from EMM2-N medium to nitrogen-rich YPD medium. Before nitrogen starvation (0d), 22 days after nitrogen starvation (22d), and at indicated time points after the transfer to YPD (6h – 24h), cell samples were fixed and stained for FACS measurement of DNA content. 1C and 2C indicate peaks of cells with unreplicated and replicated DNA, respectively. **(C and D)** WT (M161) and *mis4-450* (M162) mutant cells were fixed, stained with DAPI, and examined by fluorescence microscopy after transfer to nitrogen-rich YPD medium. Representative images of cells 20 and 22 h after the transfer are shown in (C). Images of cells before the transfer (- N22d), which are presented in Fig. 2C and those after 22-day nitrogen starvation, are shown for comparison. Arrows indicate dikaryon-like cells, in which two hemispherical interphase nuclei reside in a single cell. Bar, 10μm. In (D), more than 100 binucleated cells were examined at the indicated time points after the transfer and classified into four categories, and frequencies of cells in these categories are presented as a stacked bar graph: unseptated cells with dividing mitotic nuclei (filled in black), septated cells containing nuclei in each compartment (filled in white), unseptated dikaryon-like cells (filled in dark gray), and septated dikaryon-like cells (filled in light gray).

**Figure 4.**
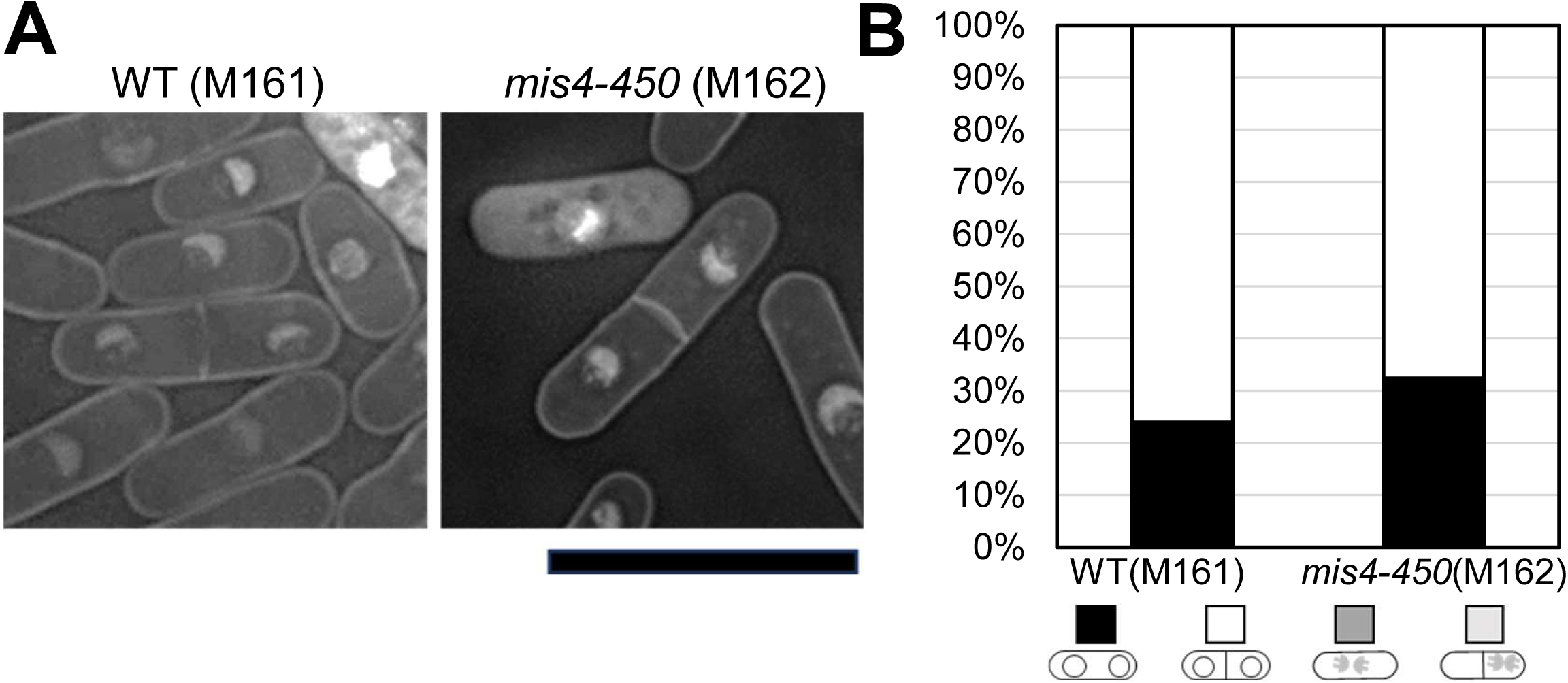
*mis4-450* mutant cells show the dikaryon-like phenotype only after exit from G0 phase. **(A and B)** WT (M161) and *mis4-450* (M162) mutant cells were continuously cultivated in nitrogen-rich YPD medium so as not to enter G0 phase, and fixed for DAPI staining. Fluorescence micrographs, which were taken and processed as described in Fig. 2C, are shown in (A). Bar, 10μm. In (B), binucleated cells were examined and classified as described in Fig. 3D. Frequencies of cells in each category are presented as a stacked bar graph. Dikaryon-like cells were not found in either WT (M161) or mutant (M162) strains. More than 80 cells were examined in each strain.

## Discussion

Mis4, an evolutionarily conserved protein in eukaryotes, functions as a cohesin loader in vegetatively proliferating cells to ensure proper cohesion of replicated sister chromatids until the onset of mitotic anaphase (Biggins and Murray, 1999; Murayama and Uhlmann, 2014). Here, we show that Mis4 has an unexpected function in non-dividing, quiescent cells, ensuring proper nuclear separation after recovery from quiescence. Although *mis4-450* (M162) mutant cells can enter G0 in response to nitrogen depletion in the medium, they fail to separate daughter nuclei when resuming mitosis upon transfer to nutrient-rich YPD medium. In the mutant, a single cell often contains two interphase-like nuclei, presumably due to insufficient separation of daughter nuclei, which are distributed equally to sister cells in the wild-type, during mitosis. This mitotic abnormality, which occurs during recovery from G0 phase in *mis4-450* mutant cells is distinct from phenotypes of *mis4* mutants previously reported (Furuya et al., 1998). When cultivated in YPD medium at a restrictive temperature (36 °C), *mis4-242 ts* mutant cells exhibit premature separation of sister chromatids after passage through S phase. Lagging and/or unequally segregated chromosomes are frequently observed during the subsequent M phase, consistent with Mis4’s function as a cohesin loader. Thus, observations in this report strongly suggest that Mis4 has an additional role in G0 phase, which may not be related to sister chromatid cohesion.

While the molecular role of Mis4 in G0 quiescent cells remains obscure, we suspect that it may be related to formation of the mitotic spindle. In a *S. pombe* strain harboring a cold-sensitive mutation in one of two α-tubulin genes (*nda2-KM52* mutant strain), two separated hemispherical nuclei, located close to each other at a position displaced from the cell center, were frequently observed in septated cells after prolonged incubation at a restrictive temperature (22 °C) (Toda et al., 1983; Umesono et al., 1983). This mitotic abnormality resembles that observed in *mis4-450* mutants. Thus, considering that *nda2* mutant cells are supposedly depauperate in functional microtubules, a major component of the mitotic spindle, due to the lack of one of two α-tubulins, Mis4’s function in non-dividing quiescent cells may somehow ensure proper formation of the mitotic spindle upon recovery from quiescence. Another possibility is that Mis4 may be involved in the septation initiation signaling network (SIN). Cells with two interphase nuclei in *mis4-450* mutants are similar to *S. pombe* dikaryon cells (Okazaki and Niwa, 2008). They showed that transient inactivation of the SIN pathway using a *ts cdc7* mutant, which prevents septum formation and subsequent cytokinesis after nuclear division, resulted in formation of dikaryon cells in *S. pombe*. Thus, in G0 phase, Mis4 may regulate timing of septum formation in the first mitotic cell cycle after recovery.

Unlike some *shk* mutants previously reported (Sajiki et al., 2009), such as *wis1* and *sty1*/*spc1* mutants, which are defective in the MAP kinase signaling pathway and which exhibit greatly reduced viability 1 day after transfer to G0-inducing EMM2-N medium, *mis4-450* mutant cells lose viability only after prolonged (>15 days) incubation in EMM2-N medium, suggesting that Mis4 is required for maintenance of G0 quiescence. Importantly, defects caused by the *mis4-450* mutation in G0 phase cannot be compensated once mitotic cell cycle progression resumes, as mutant cells lost viability and exhibit the insufficient nuclear separation phenotype during incubation in YPD medium at 26 °C, conditions in which Mis4 protein with *mis4-450* mutations (L296F and P368L) is supposedly functional. *mis4-450* mutant cells maintain high viability and proliferate vigorously at 26 °C as long as they do not enter G0 phase.

Maintaining wild-type cells in G0 in EMM2-N medium for long periods (> 15 days) induces various cytological changes in addition to rounding of the cell shape (Su et al, 1996). In cells incubated in EMM2-N medium for prolonged periods, enlarged vacuoles and lipid droplets occupy most of the cytoplasm, whereas the nucleus becomes flat and compact near the cell periphery. Mis4 may modulate chromatin structure in these nuclei so as to regulate expression of genes involved in maintenance of the functional SPB and/or proper microtubule structure that are essential for the spindle formation. In G1 phase, Mis4 is required to maintain proper expression levels of genes specifically in subtelomeric regions, to which Swi6/HP1 protein and cohesin bind interdependently (Dheur et al., 2011). Swi6/HP1 binds to histone H3, the 9^th^ lysine residue (K9) of which is methylated (Grewal and Jia, 2007), and fission yeast heterochromatin proteins, including Clr4 methyltransferase, which methylates the K9 residue of histone H3, ensures cell survival in G0 phase by methylating histone H3K9 in euchromatic regions, consequently, reprogramming the gene transcription profile (Joh et al., 2016). Mis4 may thus interact with heterochromatin proteins directly or indirectly, to establish the G0-specific gene transcription profile.

In summary, here we report a novel role of *S. pombe* Mis4 in G0 quiescent cells. Mis4’s function in G0, which is apparently distinct from that for sister chromatid cohesion in proliferating cells, is required for full separation of daughter nuclei during subsequent mitosis in cells resuming cell cycle progression, and for cell viability. Considering that Mis4/NIPBL is evolutionarily conserved from yeasts to humans and that most human cells are in the quiescent state, our findings may facilitate development of novel therapies for human diseases caused by Mis4/NIPBL deficiency.

## Materials and Methods

### Strains and Media

*S. pombe* wild-type haploid strain (972 *h*^-^) and its derivatives were used in this study. Isolation and gene identification of the original *mis4-450* mutant strain was reported previously (Sajiki et al., 2009). YPD (Yeast extract, Polypeptone, and Dextrose) medium and EMM2 (Edinburgh Minimal Medium 2) medium are used as rich- and synthetic minimal media, respectively, for cell cultivation (Moreno et al., 1991). EMM2 medium contains 93.4 mM (0.5%) ammonium chloride normally. Unless otherwise stated, cells were cultivated at 26 °C. Strains used in this study, M161 – M167 were constructed as follows: DNA fragments containing the ORF of the wild-type or a mutant version of the *mis4* gene and the hygromycin-resistant marker gene (hygR) were ligated into the pBluescript vector. Then, ∼500bp of the 3’UTR sequence of the *mis4* gene were ligated downstream of hygR, and the terminator sequence of the *S. cerevisiae ADH1* gene was inserted between the *mis4* ORF and the HygR marker gene. DNA fragments that contain the *mis4* ORF, the *ADH1* terminator, HygR, and the *mis4* 3’UTR, were excised from the vector plasmid and introduced to wild-type cells by transformation (**Fig. 1B**). Hygromycin-resistant strains were selected on YPD-solid medium containing 500 μg/mL of hygromycin, and replacement of the genomic *mis4* gene with the introduced allele was confirmed by sequencing.

### Nitrogen starvation, cell number and viability measurements, and flow cytometry

The procedure for nitrogen starvation has been described (Sajiki et al., 2009; Su et al., 1996). Briefly, cells grown in EMM2 liquid medium at a concentration of ∼2 x 10^6^ cells were harvested and resuspended in EMM2 medium lacking ammonium chloride (EMM2-N) after washing twice in EMM2-N. Cells were then cultured with shaking. Cell number was measured using a Beckman Coulter Z2 Cell and Particle Counter. To measure DNA content by flowcytometry, cells fixed in 70% ethanol were washed with 50 mM sodium citrate (pH 7.5), and stained with 1 μM SYTOX Green (Thermo Fisher Scientific) after RNase A treatment (0.1 mg/mL, 37 °C for 2 h) and Proteinase K digestion (1 mg/mL, 50 °C for 1 h) (Morimoto et al., 2022). Fluorescence intensity of cells was then measured using a FACS Calibur (BD Biosciences). Cell viability was expressed as the ratio of colonies formed on YPD solid medium to the total number (ca. 1000) of cell bodies plated after 4-day incubation at 26 °C.

### Light microscopy and image processing

For visualization of nuclear DNA, cells were fixed with 2% glutaraldehyde for 30 min on ice, washed three times with phosphate buffered saline (PBS), and then the suspension of fixed cells was mixed with an equal volume of DAPI (4’,6-diamidino-2-phenylindole, 50 μg/mL) solution on a microscope slide. DAPI-stained cells were observed using Delta Vision system (Applied Precision) equipped with a 100x oil-immersion lens (Plan Apo, NA=1.4, Olympus). Z-section images were serially acquired at intervals of 0.2 μm. Projection of serial-sectioned images and deconvolution were performed using softWoRx software (version 6.5.2, AIRIX).

## Acknowledgements

We thank Fumie Masuda and Saeko Soejima for technical assistance, Dr. Yusuke Toyoda for critical reading of the manuscript and lab members of OIST G0 cell unit for helpful discussions. Generous support from the Okinawa Institute of Science and Technology Graduate University and from Kurume University is gratefully acknowledged. This work was supported by Grants-in-Aid for Scientific Research(C) from the Japan Society for the Promotion of Science (20K06648 to Saitoh).

## Author Contributions

Conceptualization, SS and MY; Methodology, MS and SS; Investigation, all authors; Writing – Original Draft, MS and SS; Writing – Review & Editing, MS, SS and MY; Supervision, SS and MY; Funding acquisition, SS and MY.

---

**Supplementary Figure 1.**
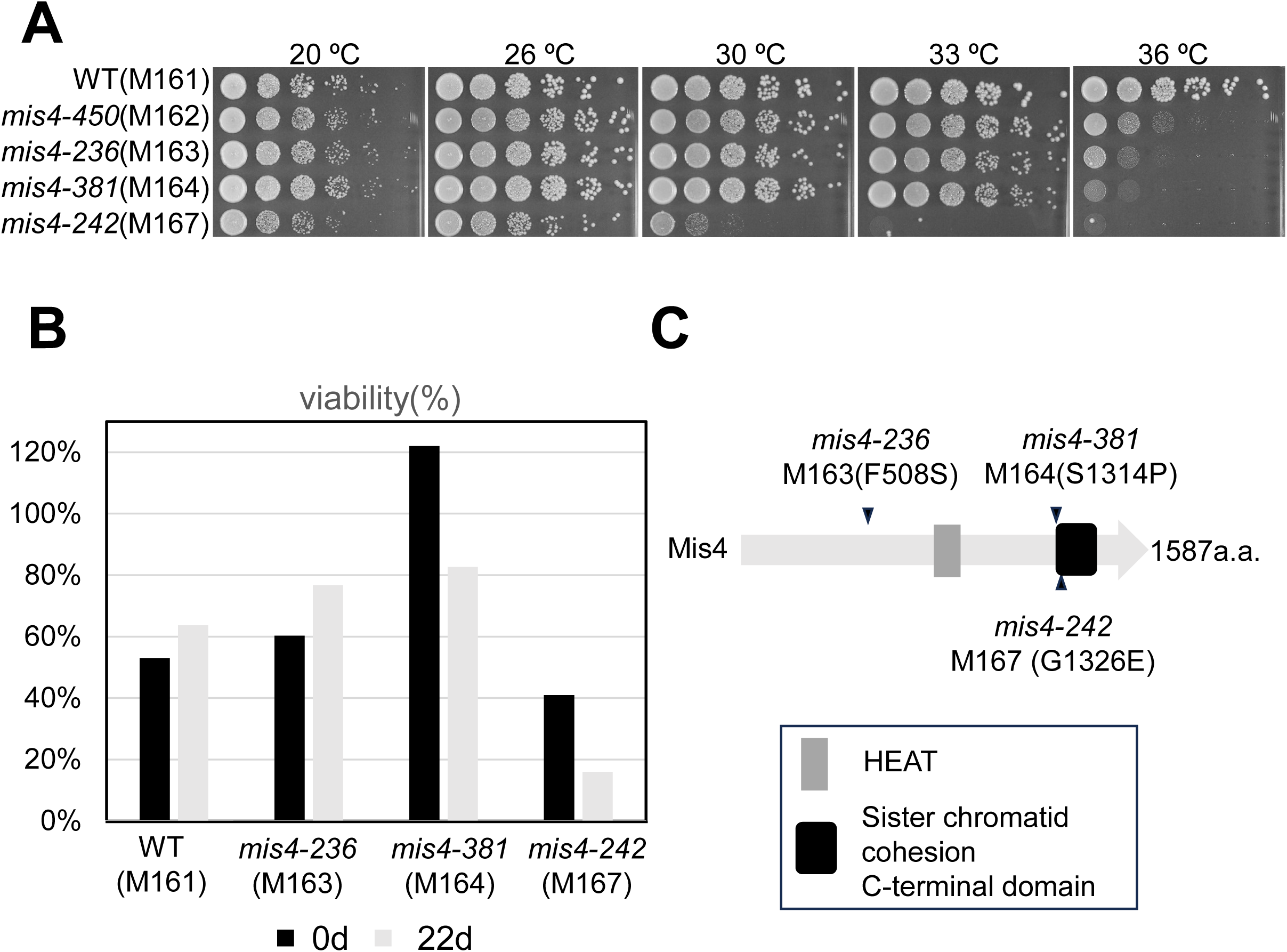
**(A)** Aliquots of 5×10^4^ cells of WT (M161), *mis4-450* (M162), *mis4-236* (M163), *mis4-381* (M164) and *mis4-242* (M167) mutant strains were diluted serially 10-fold, spotted onto YPD solid media and incubated at the indicated temperatures. Media were incubated at 20℃ for 7 days, at 26℃ for 4 days, and above 30℃ for 3 days. In M167, M163, and M164 strains, point mutations corresponding to *mis4-242* (G1326E), *mis4-236* (F508S) and *mis4-381* (S1314P) alleles were introduced to the genomic *mis4* gene of the wild-type background strain by homologous recombination. **(B)** Cell viability of the *mis4-236* (M163), *mis4-381* (M164) and *mis4-242* (M167) mutant strains were measured before (filled in black) and after (filled in gray) 22-day nitrogen starvation. **(C)** Positions of missense mutations in the *mis4-236*, *-242* and *-381* are schematically shown with characteristic domains in Mis4/Scc2/Nipped-B family proteins. See the legend of Fig. 1B for domain details.

